# GrgA controls *Chlamydia trachomatis* growth and development by regulating expression of transcription factors Euo and HrcA

**DOI:** 10.1101/2020.07.08.194431

**Authors:** Wurihan Wurihan, Yi Zou, Alec M. Weber, Korri Weldon, Yehong Huang, Zheng Gong, Zhongzi Lou, Samantha Sun, Chengsheng Zhu, Xiang Wu, Jizhang Zhou, Yaqun Wang, Zhao Lai, Huizhou Fan

## Abstract

The obligate intracellular bacterium *Chlamydia trachomatis* is an important human pathogen whose biphasic developmental cycle consists of an infectious elementary body and a replicative reticulate body. Whereas σ^66^, the primary sigma factor, is necessary for transcription of most chlamydial genes throughout the developmental cycle, σ^28^ is required for expression of some late genes. We previously showed that the *Chlamydia-*specific transcription factor GrgA physically interacts with both of these sigma factors and activates transcription from σ^66^- and σ^28^-dependent promoters *in vitro*. Here, we investigate the organismal functions of GrgA. We show that GrgA overexpression decreased RB proliferation via time-dependent transcriptomic changes. Significantly, σ^66^-dependent genes that code for two important transcription repressors are among the direct targets of GrgA. One of these repressors is Euo, which prevents the expression of late genes during early phases. The other is HrcA, which regulates gene expression in response to heat shock. The direct regulon of GrgA also includes a σ^28^-dependent gene that codes for the putative virulence factor PmpI. Conditional overexpression of Euo and HrcA also inhibited chlamydial growth and affected GrgA expression. Transcriptomic studies suggest that GrgA, Euo, and HrcA have distinct but overlapping indirect regulons. Furthermore, overexpression of either GrgA leads to decreased expression of numerous tRNAs. These findings indicate that a GrgA-mediated transcriptional regulatory network controls *C. trachomatis* growth and development.

**IMPORTANCE:** *Chlamydia trachomatis* is the most prevalent sexually transmitted bacterial pathogen worldwide and is a leading cause of preventable blindness in under-developed areas as well as developed countries. Previous studies showed that the novel transcription factor GrgA activated chlamydial gene transcription *in vitro*, but did not addressed the organismal function of GrgA. Here, we demonstrate growth inhibition in *C. trachomatis* engineered to conditionally overexpress GrgA. GrgA overexpression immediately increases the expression of two other critical transcription factors (Euo and HrcA) and a candidate virulence factor (PmpI), among several other genes. We also reveal chlamydial growth reduction and transcriptomic changes including decreased GrgA mRNA levels in response to either Euo or HrcA overexpression. Thus, the transcription network controlled by GrgA likely plays a crucial role in chlamydial growth and pathogenesis.

## INTRODUCTION

The Centers for Disease Control and Prevention (CDC) reports that chlamydia is the most common notifiable disease in the United States. Caused by infection with *Chlamydia trachomatis*, this sexually transmitted disease (STD) has comprised the majority of all STDs reported to CDC since 1994 (1). The World Health Organization estimates 131 million new cases of *C. trachomatis* infection occur annually world-wide (2). Although infection with *C. trachomatis* is usually asymptomatic, untreated chlamydial infection is associated with high rates of infertility, pelvic inflammatory syndrome, abortion and/or premature birth, and ectopic pregnancy (1, 2). These serious complications exemplify the primary burden of disease. Furthermore, three *C. trachomatis* serotypes are known to cause ocular infection and blinding trachomatous trichiasis. These clinical manifestations are still common not only in many underdeveloped countries, but also in developed nations (3).

Chlamydiae are obligate intracellular Gram-negative bacteria with a unique developmental cycle characterized by two cellular forms (4). The small, electron-dense form termed elementary body (EB) is capable of extracellular survival but incapable of proliferation. After binding to receptors on the target cell membrane, the EB is taken into a cell membrane-derived vacuole through endocytosis (5). Within the vacuole termed inclusion, the EB differentiates into a larger, less electron-dense form termed reticulate body (RB) within several hours. The RB replicates with a doubling time of 2 to 3 h. Around 24 h, some RBs start to re-differentiate back into EBs while others continue to proliferate. At the end of the cycle, EBs and residual RBs are released from host cells through either cell lysis or extrusion of entire inclusions (6).

The small *C. trachomatis* genome consists of a 1 million bp chromosome and a 7.5 kb plasmid. The chromosome carries less than 900 total protein-encoding genes and noncoding RNA genes. The plasmid encodes only 8 proteins (7). Previous cDNA microarray studies (8, 9) enumerate four successive stages of the developmental cycle. The immediate early stage is the first h when EBs are inside nascent inclusions near the plasma membrane. A small number of crucial genes are transcribed in this stage to establish an intracellular niche that enables EB survival, development into RBs, and eventual delivery of the inclusion to a perinuclear region. During the subsequent early stage, an additional number of genes are transcribed to complete the conversion of EBs into RBs. Midcycle commences upon the completion of EB-to-RB conversion and ends when RBs starts to differentiate back into EBs. Almost all genes are transcribed during this stage. Lastly, transcription of a smaller set of genes is initiated and/or upregulated before and during the late stage.

In bacteria, all genes are transcribed by one type of RNA polymerase (RNAP). The RNAP holoenzyme consists of a catalytic core enzyme with one of several different sigma factors (σs) that recognize various promoters (10). *C. trachomatis* encodes three σs. A super majority of *C. trachomatis* promoters are σ^66^-dependent, while some late genes possess σ^28^ promoters or both σ^66^ and σ^28^ promoters (11–13). To date, only two late genes are thought to carry a σ^54^ promoter (14). Consistent with their roles in the developmental cycle, expression of the three σs is also temporally regulated (8, 15). σ^66^ mRNA is detected on microarray as early as 3 h postinoculation (hpi), whereas σ^28^ and σ^54^ mRNAs are not detected until 8 hpi (8).

Transcription activities of the RNAP are regulated by transcription factors (TFs). Interestingly, *C. trachomatis* encodes fewer than 20 TFs despite a complicated developmental cycle (16). Most of these TFs regulate gene expression in response to nutrient and mineral availability (17–30). In contrast, the transcription repressor HrcA controls response to heat shock (31, 32).

Only two *C. trachomatis* transcription factors demonstrate ability to control the chlamydial developmental cycle, Euo and CtcC. Euo is produced immediately after EBs enter host cells (8, 33, 34) and binds late gene promoters to suppress transcription (11–13). CtcC is a part of a two-component system and is predicted to function as an activator of the σ^54^-RNAP holoenzyme on the basis of orthologs in other bacteria (25).

GrgA is the newest chlamydial TF. Identified via promoter DNA pulldown, GrgA physically interacts with σ^66^ and σ^28^, and activates transcription from both σ^66^- and σ^28^-dependent promoters *in vitro* (35–37). In this work, we investigated the organismal functions of GrgA. Through overexpression, growth characterization, transcriptomic studies, and protein expression analyses, we identify a GrgA-directed transcriptional regulatory network (TRN) that likely plays a critical role in chlamydial growth and development.

## RESULTS

### GrgA overexpression inhibits *Chlamydia trachomatis* growth

In our initial attempt to overexpress GrgA, we placed the GrgA open reading frame (ORF) downstream of a *Neisseria meningitidis* promoter (P_nm_) in the pGFP::SW2 plasmid (38) (Fig. S1A). With this resultant pGFP-CmR-GrgA::SW2 vector (Fig. S1B), we failed to obtain transformants after three independent attempts despite consistent transformant production by the control pGFP::SWP plasmid. These negative data were an early suggestion that GrgA overexpression may be toxic.

Next, we constructed the pTRL2-NH-GrgA vector by placing His-tagged GrgA downstream of a P_tet_ promoter (Fig. S1C). pTRL2-NH-GrgA transformants of CtL2 were readily appreciable following two passages of selection with penicillin. These uncloned transformants formed a similar number of notably smaller inclusions after ATC treatment (Fig. S2). We proceeded by generating clonal populations, of which one was subject to Western blotting to confirmed successful overexpression after ATC treatment. Both anti-GrgA and anti-His-tag antibodies were able to individually detect recombinant His-tagged GrgA (Fig. S3).

To further characterize the apparent effects of GrgA overexpression on chlamydial growth and development, we initiated ATC treatment of the clonal population analyzed in Fig. S3 at one of four time points: 0, 8, 18 and 24 h post-inoculation (hpi). We harvested one set of cultures at 30 hpi and quantified their yield of progeny EBs. Images were acquired for another set of cultures at 36 hpi. The control vector-transformed CtL2 showed no difference in the EB production between the non-induce and induced cultures (Fig. 1A). This finding was corroborated by a lack of difference in the number, size, and RFP intensity of inclusions (Figs. 1B, S4A-C). The GrgA-transformed CtL2 produced a statistically significant lower number of EBs when ATC induction occurred between 0 and 18 hpi (Fig. 1C). Although a decrease in EB yield was not observed when ATC induction was conducted at 24 hpi (Fig. 1C), direct imaging of all conditions at 35 hpi did reveal reduced inclusion area and RFP intensity for the 24 hpi condition (Figs. 1D, S4D-F). These findings indicate that delicate regulation of physiological GrgA concentrations during the first 18 h is critical for adequate CtL2 development and growth.

**Fig. 1.**
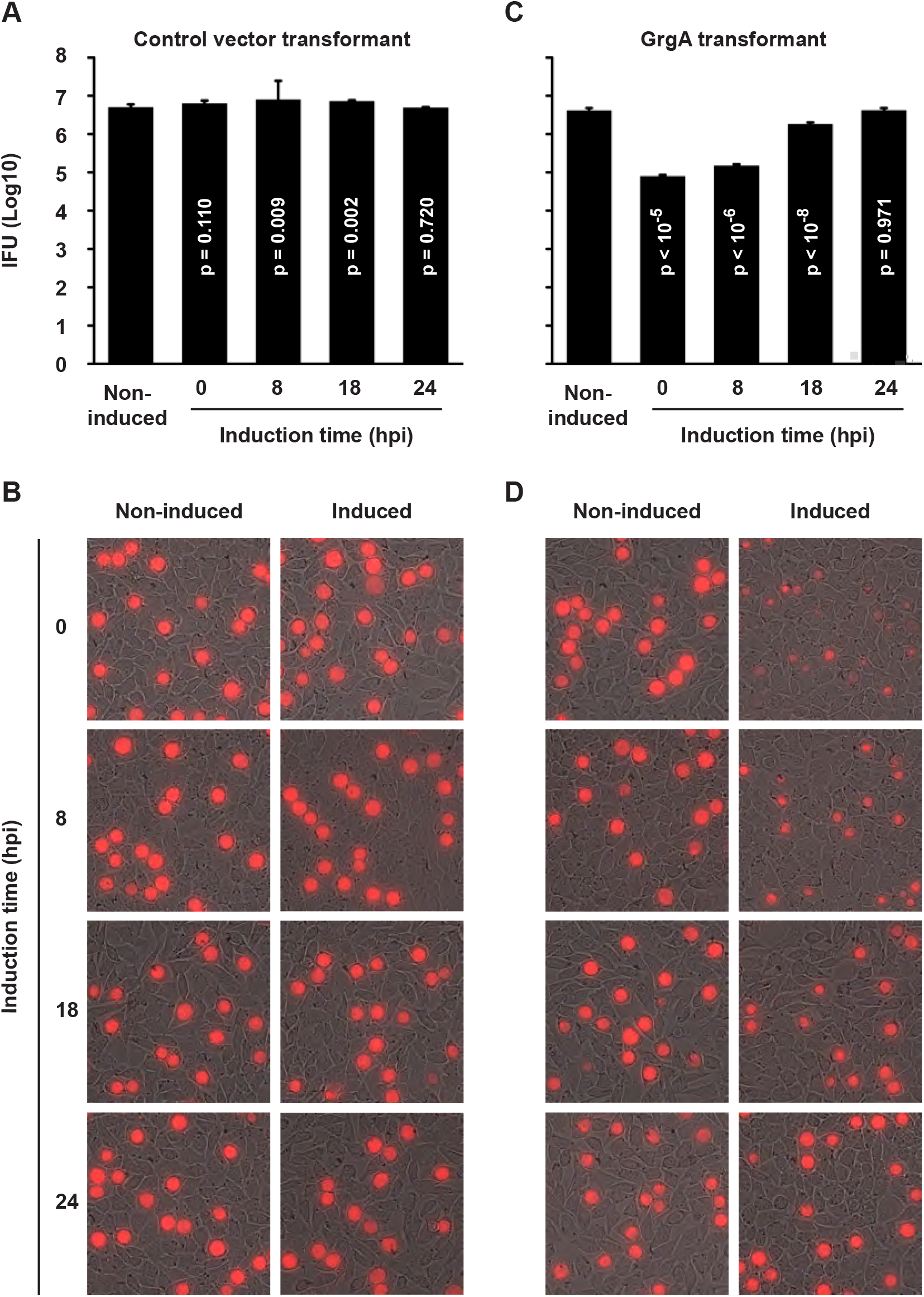
Ectopic expression of GrgA inhibits chlamydial growth. ATC was added to cultures of control vector transformants (A, B) and GrgA transformants (C, D) at the indicated h post-inoculation (hpi). Number of progeny EBs formed were determined at 30 hpi (A, C); RFP-expressing inclusions in live cultures were imaged at 35 hpi (B, D). (A, C) Data are averages ± standard deviations of triplicate experiments. See Fig. S4 for number of inclusions per image, inclusion sizes, and inclusion RFP intensities associated with panels B and D.

### Deletion of σ^66^-binding domain from GrgA fully eliminates overexpression-induced inhibition while deletion of σ^28^-binding domain only partially reverses

Our previous *in vitro* studies showed that GrgA activates both σ^66^-dependent and σ^28^-dependent transcription. GrgA binds σ^66^ and σ^28^ at residues 1-64 and 138-165, respectively (35, 36) (Fig. 2A). We constructed GrgA expression vectors that lack these regions to understand how each σ factor might contribute to chlamydial inhibition following GrgA overexpression, if at all. Expression of these GrgA deletion mutants in clonal populations of transformants was then detected by western blotting (Fig. S5). In contrast to full-length GrgA overexpression (Figs. 1, S4), Δ1-64 GrgA overexpression showed no adverse effects on chlamydial growth (Figs. 2B-C, S6A-C). Δ138-165 GrgA overexpression reduced progeny EB production when ATC was added at 0, 8, and 18 hpi, albeit by magnitudes about 10-fold less than what was previously observed after full-length GrgA overexpression (Fig. 2D-E). Δ138-165 GrgA overexpression decreased the inclusion size and RFP intensity only at 0 hpi (Figs. 2D-E, S6D-F). These findings indicate that interaction with σ^66^ is absolutely required for GrgA overexpression-induced inhibition, whereas interaction with σ^28^ also plays a significant role.

**Fig. 2.**
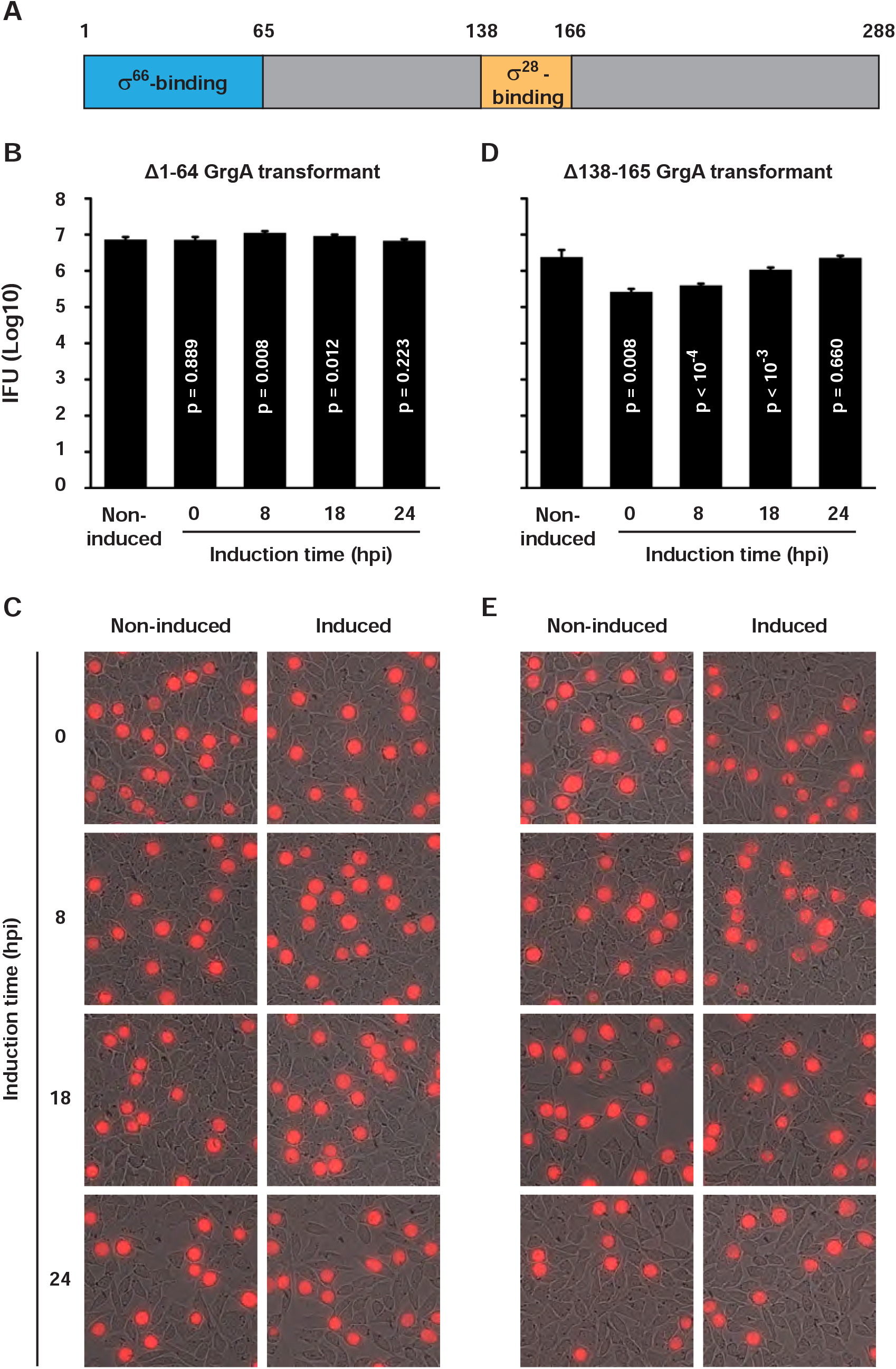
Differential effects of σ^66^-binding-defective Δ1-64 GrgA and σ^28^-binding-defective Δ138-165 GrgA overexpression on chlamydial growth. Domain structure of the GrgA protein is shown in (A). Data were acquired in the same manner as for Fig. 1. See Fig. S6 for number of inclusions per image, inclusion sizes, and inclusion RFP intensities associated with panels C and E.

### GrgA overexpression inhibits RB replication and volume expansion in a σ^66^-binding domain-dependent manner

We further performed TEM for ATC-treated GrgA transformants from 8 to 14 hpi. This analysis revealed a statistically significant 16% decrease in RB size in the ATC-treated cultures (Figs. 4A, 4B). This finding implies that GrgA overexpression not only impacts the EB-to-RB differentiation process, but also impedes RB volume expansion after division.

We employed quantitative confocal microscopy to investigate the effect of GrgA overexpression on RB proliferation. In cells infected with control vector transformants, ATC treatment did not affect the number of RBs per inclusion. In cells infected with GrgA transformants, ATC treatment during the period of 8 to 14 hpi caused a 50% reduction in the number of RBs per inclusion when compared to non-treated control cultures (Fig. 3C, D). Successive quantitative PCR (qPCR) analysis was conducted to corroborate these RB enumeration data. In GrgA transformants, a significant reduction in genome copy number was readily detected 2 h after ATC induction, which became progressively severe in following hours (Fig. 3E). Together, we infer from the findings presented in Fig. 3 that RB proliferation is inhibited by GrgA overexpression. Furthermore, confocal microscopy analyses (Fig. S7) of GrgA deletion mutants also demonstrate that the inhibition of RB proliferation is highly dependent on the ability of GrgA to interact with σ^66^, and less dependent on its ability to interact with σ^28^. These results are consistent with cellular growth data presented in Figs. 2, S6.

**Fig. 3.**
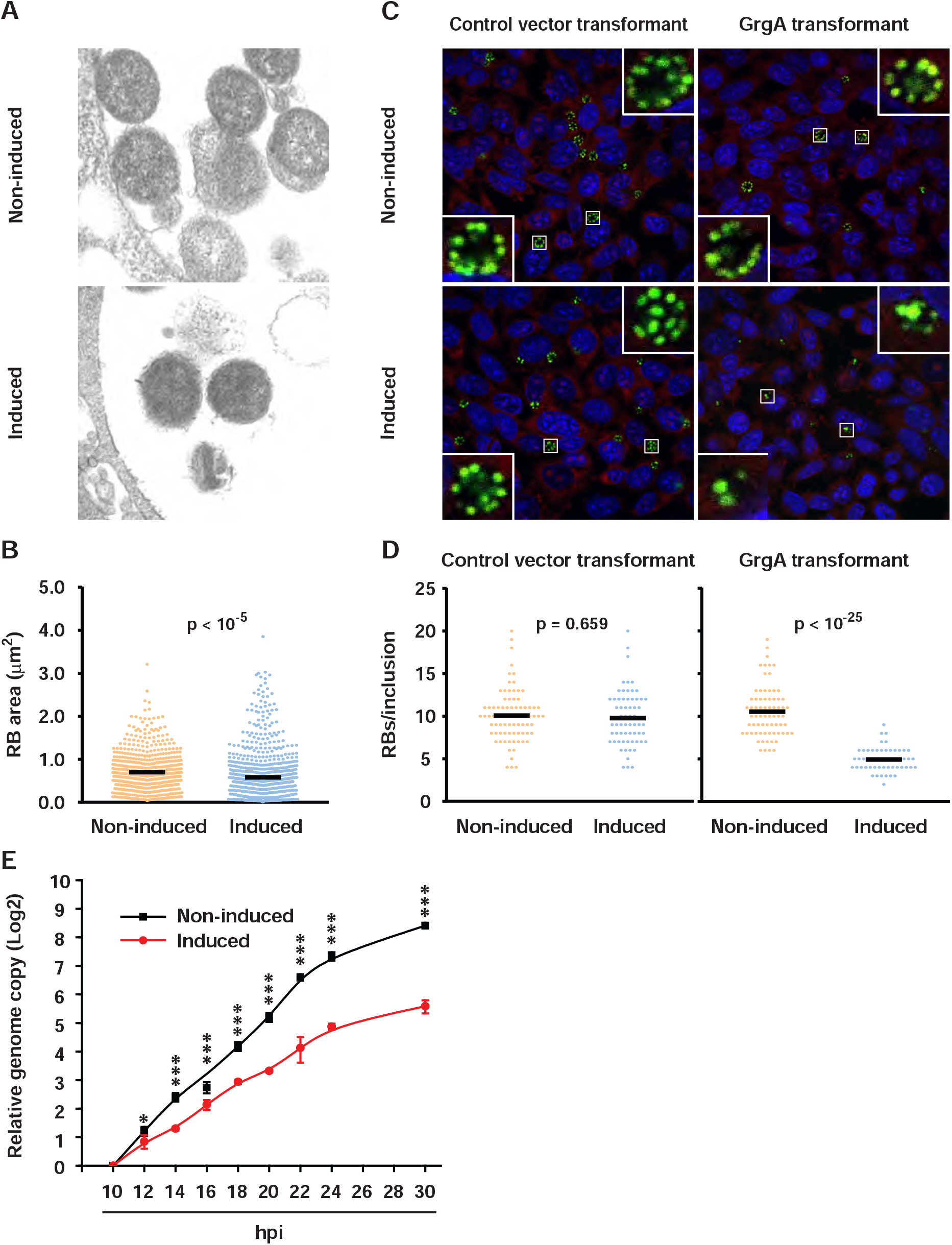
GrgA overexpression decreases RB volume expansion and replication. (A) Representative TEM images of RBs formed by GrgA transformants in infected cells in either the absence or presence of ATC between 8 and 14 hpi. (B) Scattergram of the RB area data obtained from TEM images. (C) Confocal microscopy images of control and GrgA transfor-mants in cells cultured with or without ATC during the periods of 8 to 14 hpi. (D) Scattergram of the number of RBs per inclusion analyzed by confocal microscopy. (E) Relative genome copy numbers in GrgA transformants cultured with or without ATC at 10 hpi. Data are averages +/− standard deviations of triplicate experiments. *, p < 0.05; ***, p < 0.001.

### GrgA overexpression-mediated global transcriptomic changes include upregulated Euo, HrcA, and PmpI expression and decreased tRNA expression

We performed RNA-seq analyses to determine the molecular mechanism underlying GrgA overexpression-induced growth inhibition. Since few chlamydial RNA-seq studies with unpurified organisms existed in the literature at the time, our pilot RNA-seq experiments were conducted to optimize the timing of ATC induction and sample harvesting. GrgA transformants were treated with or without ATC within two time periods: 12 to 16 hpi and 17 to 21 hpi (Table S1, S2). As expected, the mapping rates of samples prepared at 16 hpi were more than 3-fold lower than the rates of those prepared at 21 hpi (Table S3). With the exception of rRNAs, which were depleted prior to library preparation, RNAs of all chlamydial genes could be detected at 16 hpi despite this notable decrease. In both sets of experiments, mRNA reads of two transcription repressors Euo and HrcA were noticeably increased in ATC-induced cultures. For the 12 to 16 hpi induction, Euo and HrcA reads increased by 3.1 and 2.8 fold, respectively. For the 17 to 21 hpi induction, they increased by 3.1 and 1.9 fold, respectively.

Subsequent RNA-seq studies were conducted with samples harvested at 16 hpi. This time point corresponds to the mid-log phase of RB replication, whose regulatory mechanisms are most interesting to us. We repeated RNA-seq analyses with ATC-treated biological replicates for the 12 to 16 hpi time period to generate statistic power. Consistent with our previous two RNA-seq studies, mRNA reads of both Euo and HrcA increased by a statistically significant level in response to ATC treatment. Euo increased by 3.3-fold, second only to the ATC-induced increase in GrgA mRNA reads. HrcA increased 2.1-fold, the 5th largest increase. RNA reads of 89 other genes also increased significantly (i.e., *P* < 0.05), whereas those of the remaining 86 genes were significantly decreased (Table S4). Of the 86 genes with significantly downregulated RNAs, 33 were tRNA genes. Only 4 of the 37 tRNAs were not significantly downregulated (Table S5). Retrospective analyses showed that numerous tRNAs were also downregulated in previous RNA-seq studies (31 tRNAs in the experiment with ATC treatment from 12 to 16 hpi; 8 tRNAs in the experiment with ATC treatment from 17 to 21 hpi).

### Activation of *euo* and *hrcA* but not *pmpI* depends on σ^66^-binding of GrgA

To determine the contribution of GrgA overexpression-induced transcriptomic changes to chlamydial growth defects in GrgA transformants, we performed RNA-seq for CtL2 transformants of Δ1-64 GrgA and Δ138-165 GrgA with and without ATC treatment between 12 and 16 hpi (Tables S6, S7). In ATC-treated Δ1-64 GrgA transformants, induction of only a single gene was statistically significant while repression of four genes was statistically significant. In ATC-treated Δ138-165 GrgA transformants, the numbers of activated and repressed genes were both higher than those of ATC-treated full-length GrgA transformants. The sole gene activated by Δ1-64 GrgA overexpression was PmpI, which was also activated by overexpression of both full-length GrgA and Δ138-165 GrgA. Noticeably, PmpI mRNA increased after full-length GrgA overexpression (253%) and Δ1-64 GrgA overexpression (270%) by almost the same magnitude. These increases were significantly higher than the 40% increase observed after Δ138-165 GrgA overexpression.

55 genes were induced by both full-length GrgA overexpression and Δ138-165 GrgA overexpression (Fig. 4A, Table S8). 2 of the 4 genes repressed by Δ1-64 GrgA were also repressed by both full-length GrgA and Δ138-165 GrgA overexpression. The third Δ1-64 GrgA-repressed gene was also repressed by full-length GrgA. In total, 45 genes were induced by both full-length GrgA overexpression and Δ138-165 GrgA overexpression (Fig. 4B). Of these 45 genes, 28 encode tRNAs (Table S9).

**Fig. 4.**
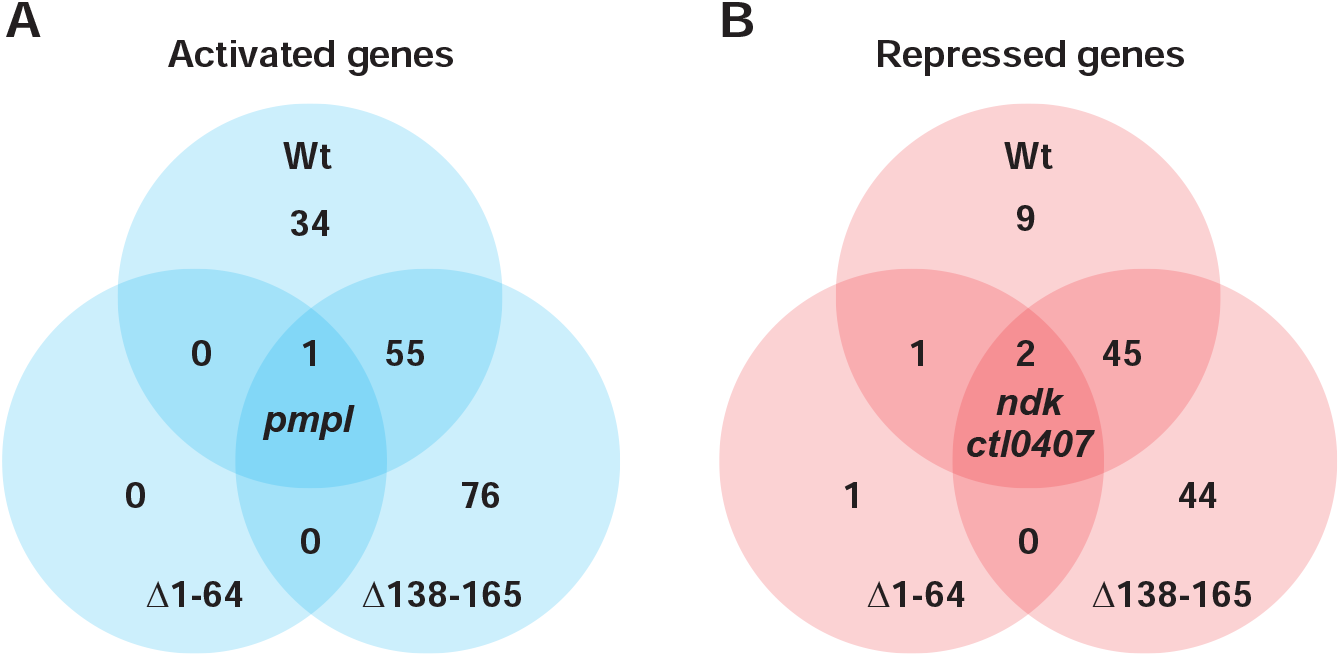
Venn diagram showing up- and down-regulated genes detected in full length (Wt) GrgA, Δ1-64 GrgA, and Δ138-165 GrgA transformants treated with ATC. The figure was prepared with data presented in Tables S8, S9. Up-or down-regulation was defined with ≥33% change with a P value ≤ than 0.05.

Taken together, comparative transcriptomic analyses suggest that nearly all transcriptomic changes (including activation of *euo* and *hrcA* but *pmpI*) induced by GrgA depend on GrgA binding of σ^66^. By contrast, fewer changes depend on GrgA binding of σ^28^. However, overexpression of the σ^28^-binding defective Δ138-165 GrgA may induce additional transcriptomic changes not seen with full length GrgA overexpression.

### *euo* and *hrcA* among genes activated immediately following GrgA overexpression

To identify genes directly targeted by GrgA, we determined changes in transcriptomic kinetics by extracting RNA at 16 hpi from non-induced cultures and cultures treated with ATC for 0.5, 1 and 2 h and (see Fig. S8 for experimental design). Results of RNA-seq reads are presented in Table S10; expression levels normalized with values of the transcriptome fragments per kilobase per million reads mapped (FPKM) are presented in Table S11. The entire transcriptome can be divided into 6 groups based on changes in the expression kinetics of individual genes (Fig. 5, Table S12). Group A contains 6 genes whose mRNAs increased by 0.5 h (Fig. 5A), although the increase was statistically significant for only four of the six mRNAs. The four statistically significantly increased mRNAs were those of Euo, PmpI [a polymorphic protein in the outer membrane and putative virulence gene (39, 40)], AroC (chorismate synthase, which is involved in aromatic amino acid biosynthesis), and LplA (lipoate protein ligase A) (Fig. 5A). These 6 genes are likely primary targets of GrgA. mRNAs of 175 genes, including hrcA, increased by 1 h (Fig. 5B). These large group of genes may be secondary or indirect targets of GrgA. mRNAs of 444 genes remained relatively constant (Fig. 5C) and are therefore unlikely targets of GrgA. mRNA levels of remaining genes decreased to various degrees (Fig. 5D-F), likely in response to expression changes of the primary and/or secondary targets.

**Fig. 5.**
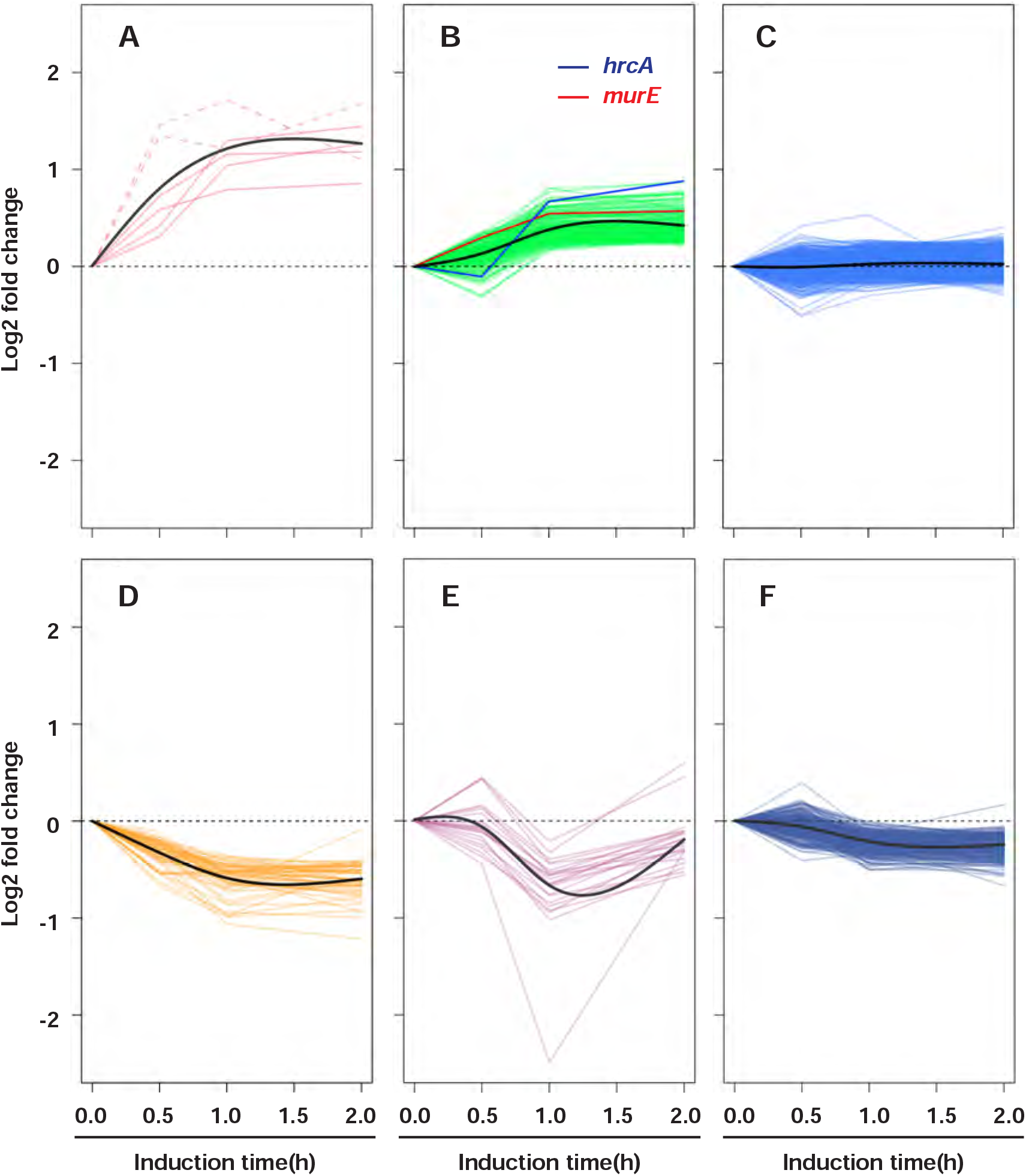
Temporal patterns of transcriptomic changes induced by GrgA overexpression. Figures were derived from RNA-sea data presented in Table S10. (A) Six early target genes were increased by 0.5 h. RNAs whose increases were statistically significant (P < 0.05) are shown in solid lines. RNAs which increased with a P value larger than 0.05 by 0.5 h are shown in dashed lines. (B) RNAs of 175 genes are stimulated by GrgA overexpression only after one hour of ATC induction. Red and blue lines are for the mRNAs of MurE and HrcA, respectively. (C) RNAs of 444 genes remained relatively constant. (D-F) Genes are down regulated following different kinetics. (A-F) Solid black lines are trend lines in respective groups.

Among the six putative “early” genes induced by GrgA (Fig. 5A, Table S12), *euo, pmpI, ctl0758* and *ctl0418* are found in single-gene units (Fig. 6A), whereas *lplA* and *aroC* in operons. Based on the genome topology (7, 41), and results of genome-wide transcription start site analyses (42), *lplA* shares a promoter with *ctl0536* (Fig. 6B), whereas *aroC* is cotranscribed with 3 other genes (Fig. 6C). Noticeably, cotranscribed mRNA reads did not increase as the reads of LplA and AroC mRNAs increased. To validate the RNA-seq data, we performed reverse transcription quantitative PCR (RT-qPCR). Among the four singly transcribed mRNAs, these analyses showed Euo and PmpI readily increased by about 2- and 3-fold, respectively, at 10 min after induction, and more than 3- and 4-fold, respectively, at 30 min (Fig. 1A). Smaller but significant increases were detected for the mRNA of *ctl0758* from 10-30 min (Fig. 6A). However, a significant (57%) increase *in ctl0418* was not detected until 30 min (Fig. 6A).

**Fig. 6.**
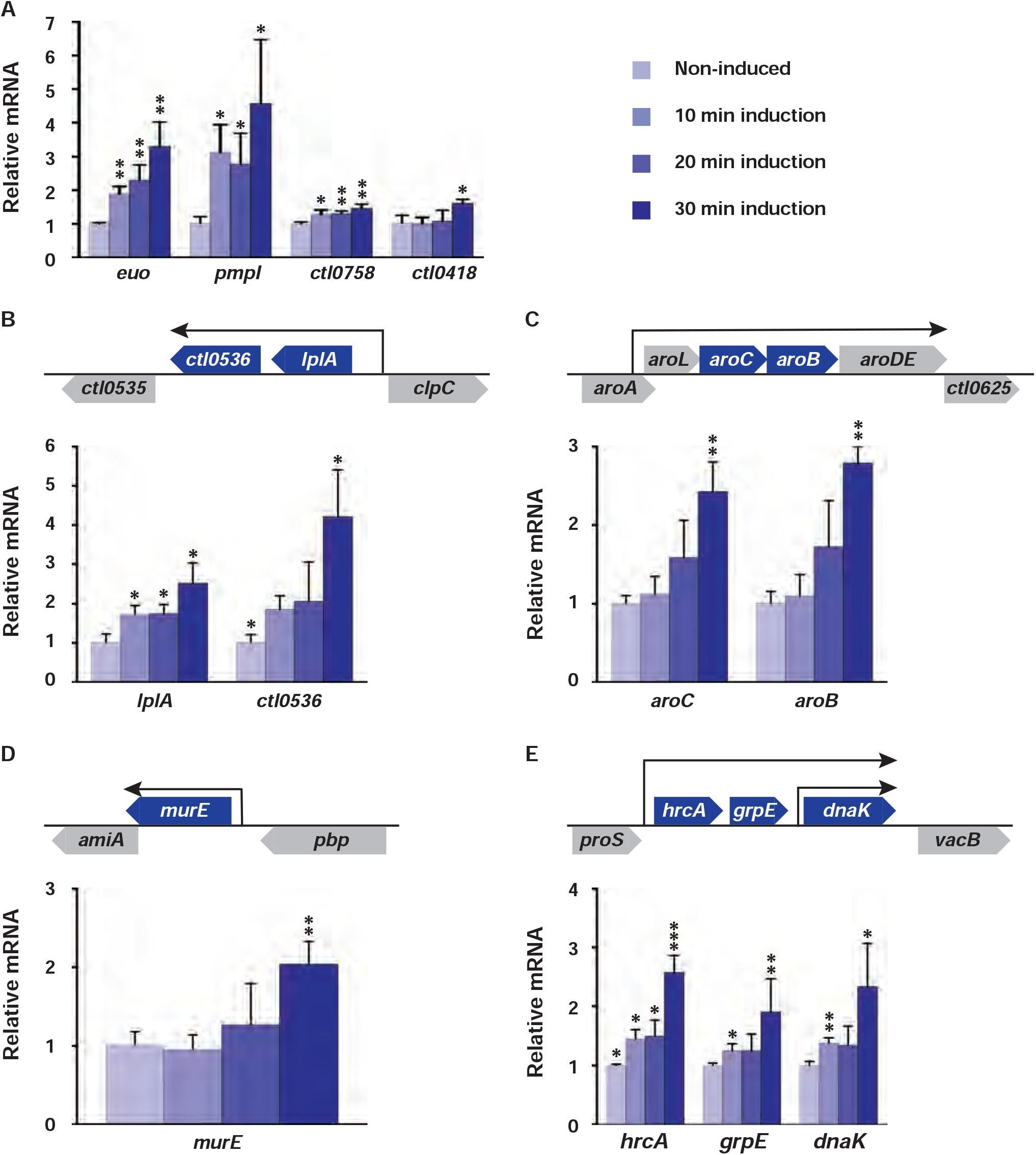
RT-qPCR detection or confirmation of early target genes of GrgA. Relative mRNA levels of 5 non-operon genes are shown in (A, D) and those in operons are shown in B, C, and E. MurE (D) and HrcA (E) were identified as early genes only by RT-qPCR. See also Fig. S12. (A-E) Data are averages ± standard deviations of triplicate experiments. *, p < 0.05; **, p < 0.01; ***, p < 0.001.

For the two operon genes (*lplA* and *aroC*), we included a transcription partner in our RT-qPCR analyses. mRNA expression trends of both LplA and its partner CTL0536 were similar to those of EUO and PmpI (Fig. 6B) with significant increases starting at 10 min. Trending increases were also found for the mRNAs of AroC and its transcription partner AroB (Fig. 6C). These results validate *lplA* and *aroC* as early genes induced by GrgA. However, failure of RNA-seq analysis to detect increased mRNAs of their transcription partners at 30 min indicate that our RNA-seq was not as sensitive as RT-qPCR.

The apparent higher detection sensitivity of RT-qPCR prompted its use to determine whether any additional genes whose mRNA reads increased at 1 h may actually be increased at 30 min as well. Our criteria for selection were 1) a read increase with *P* < 0.05, and 2) at least one fragments per kilobase per million reads mapped (FPKM) being >900 for induced samples. As an exception, because HrcA is an important TF, its mRNA was also analyzed using RT-qPCR even though the FPKM values were only 445, 329 and 334 in ATC-induced samples. RT-qPCR analysis detected apparently increased levels for all mRNAs analyzed; however, only the increases in the mRNAs of MurE (UDP-N-acetylmuramoyl-L-alanyl-D-glutamate-2,6-diaminopimelate ligase in the peptidoglycan synthesis pathway) and HrcA were statistically significant (*P* < 0.05) (Fig. S12). Further analysis confirmed that MurE mRNA increased at 30 min but not 10 or 20 min after ATC induction (Fig. 6D), but HrcA mRNA readily increased at even 10 min (Fig. 6E). *hrcA* is in an operon with two transcription partners *grpE* (which encodes heat shock protein-70 cofactor) and *dnaK* (a protein chaperone gene) although *dnaK* has an additional promoter. Similar to the HrcA mRNA, the mRNAs of GrpE and DnaK showed similar increases starting 10 min (Fig. 6E). Taken together, results presented in Fig. 6, S9 demonstrate that 5 genes (*euo, pmpI, murE, ctl0418* and *ctl0758*), which are in single-gene transcription units, and 9 additional genes in 3 operons are activated by 10 to 30 min induction of GrgA overexpression. Most likely, these 14 genes comprise GrgA’s direct regulon.

### GrgA stimulates transcription from *euo, hrcA* and *pmpI* promoters

Sequence analyses identified putative σ^66^ promoter elements in at least 5 of the 8 GrgA-regulated promoters and revealed that *pmpI* appears to carry additional σ^28^ promoter elements (Fig. S10). We constructed four transcription reporter plasmids, of which three carried a putative σ^66^ promoter of *euo, hrcA,* or *pmpI*. The remaining reporter plasmid carried the putative σ^28^ promoter of *pmpI*. Of these four constructed plasmids, all were able to successfully direct RNA synthesis except for the one carrying a putative σ^66^ promoter of *pmpI*. This suggests the cloned promoter fragment is nonfunctional. Among the other three plasmids, an increased number of transcripts were detected in the presence of GrgA (Fig. 7) indicating that GrgA activates transcription not only from the σ^66^ promoters of *euo* and *hrcA* (i.e., PdnaK2), but also from the σ^28^ promoter of *pmpI*.

**Fig. 7.**
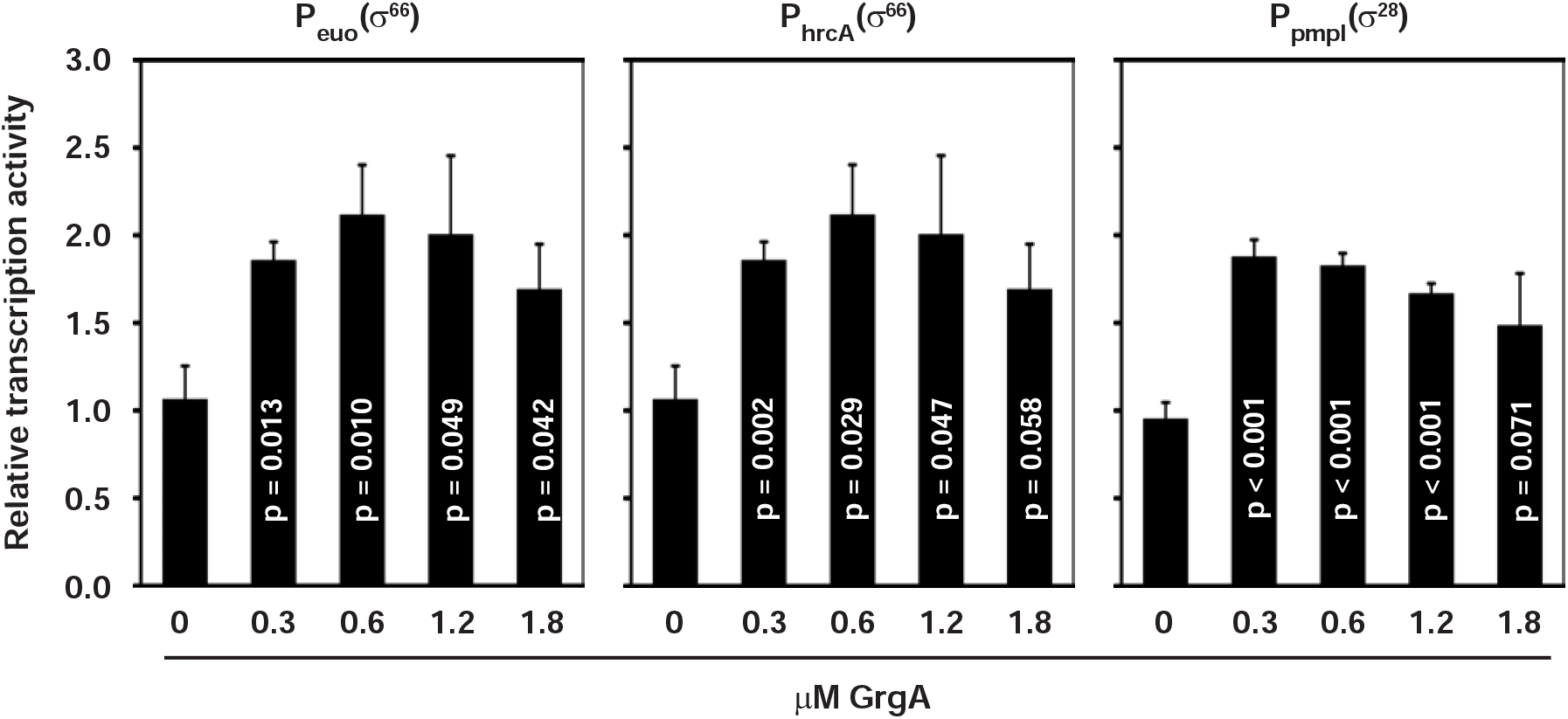
Stimulation of transcription from *euo*, *hrcA* and *pmpI* promoters by GrgA *in vitro*. Data are average ± standard deviations of triplicate experiments.

### Overexpression of either Euo or HrcA inhibits chlamydial growth

To determine the contributions of upregulated Euo and HrcA expression to GrgA overexpression-induced growth inhibition, we constructed ATC-inducible Euo and HrcA expression plasmids (Table S9) and generated CtL2 transformants thereafter. 10 nM ATC-induced overexpression of both Euo and HrcA following treatment at 0, 8, and 18 hpi caused severe to moderate growth inhibition; these effects were minimal after treatment at 24 hpi (Fig. 8). These results are similar to our earlier observations made following ATC-induced GrgA overexpression (Fig. 1). Next, we used lower ATC concentrations to induce 2-to 3-fold increases in the mRNAs of Euo and HrcA, which were comparable to their magnitudes of increase induced by ATC in GrgA transformants (Table S1, S2, S4). Noticeably, low ATC concentrations were also sufficient to cause growth inhibition (Fig. S11). These data support the notion that increased Euo and HrcA expression mediate GrgA overexpression-induced CtL2 growth.

**Fig. 8.**
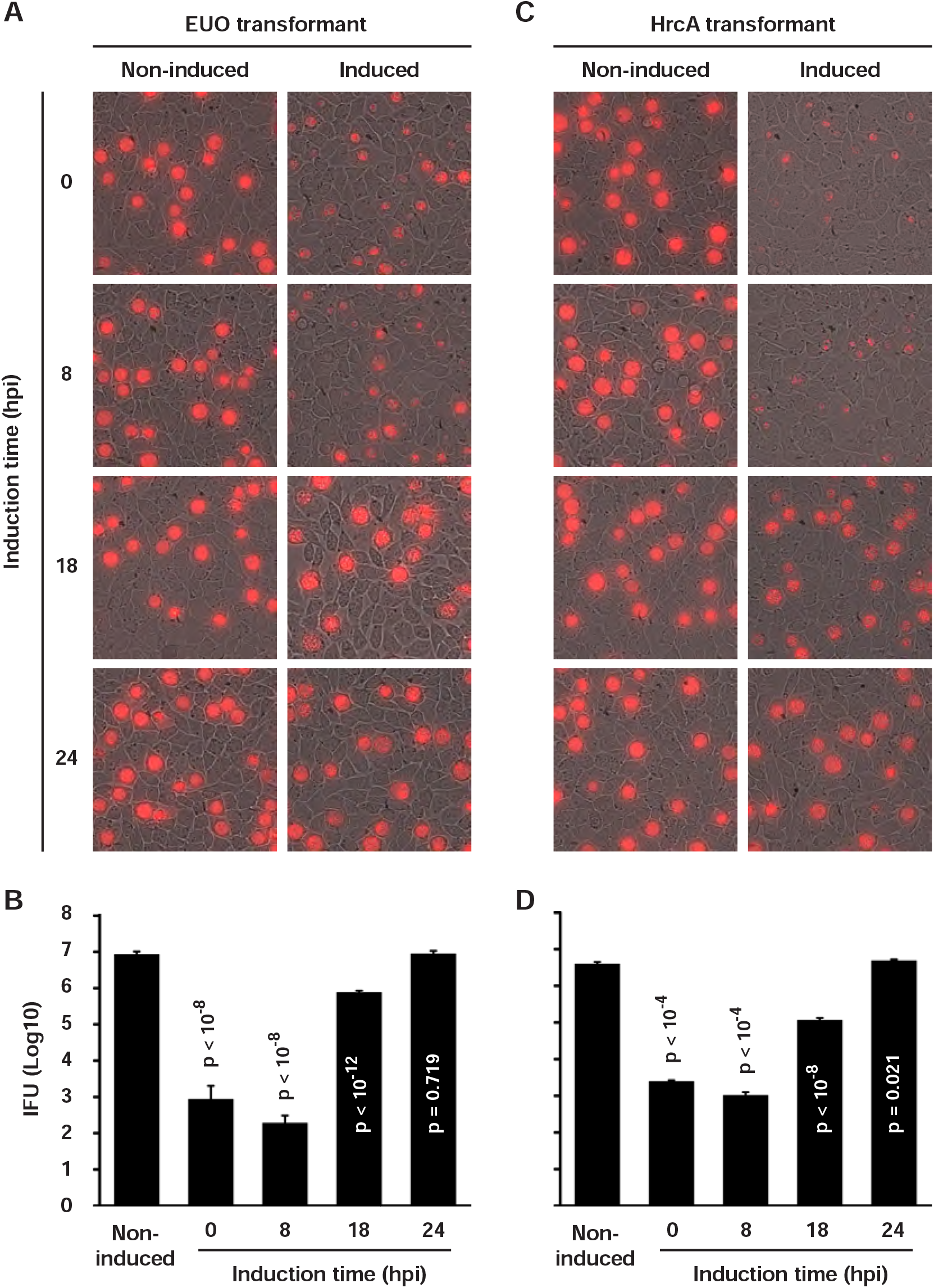
Inhibition of chlamydial growth by either Euo or HrcA overexpression. Data were acquired in the same manner as for Fig. 1.

### Identification of genes commonly regulated by GrgA, Euo and HrcA

We performed RNA-seq analyses to identify transcriptomic changes in Euo and HrcA transformants following ATC-induction between 12 and 16 hpi (Tables S13, S14). By comparing these two RNA-seq datasets with the RNA-seq dataset obtained from the GrgA transformant (Table S4), we identified the genes that are commonly regulated by GrgA, Euo, and HrcA (Fig. 9). Whereas 9 genes were commonly activated (Fig. 9A, B) in all three transformants upon ATC treatment, 11 genes were commonly repressed (Fig. 9C, D). Genes that are commonly activated or repressed in two transformants as well as in all three transformants are listed in Tables S15, S16. Noticeably, 4 of the 9 genes that are commonly activated in all three transformants encode proteins involved in DNA replication, whereas the remaining 5 commonly activated genes encode proteins with various functions (Fig. 9B). Only 6 of the 11 genes commonly repressed in all three transformants following ATC treatment encode functionally known proteins. Three (PPA, IspH and FabI) catalyze metabolic reactions while the other three either constitute a protein translocase (YajC) or serve as secretion effectors (CTL0874 CTL0887) (Fig. 9D). These commonly activated and repressed genes may serve as upstream regulators of RB growth and proliferation or their expression levels are controlled consequent to growth inhibition.

**Fig. 9.**
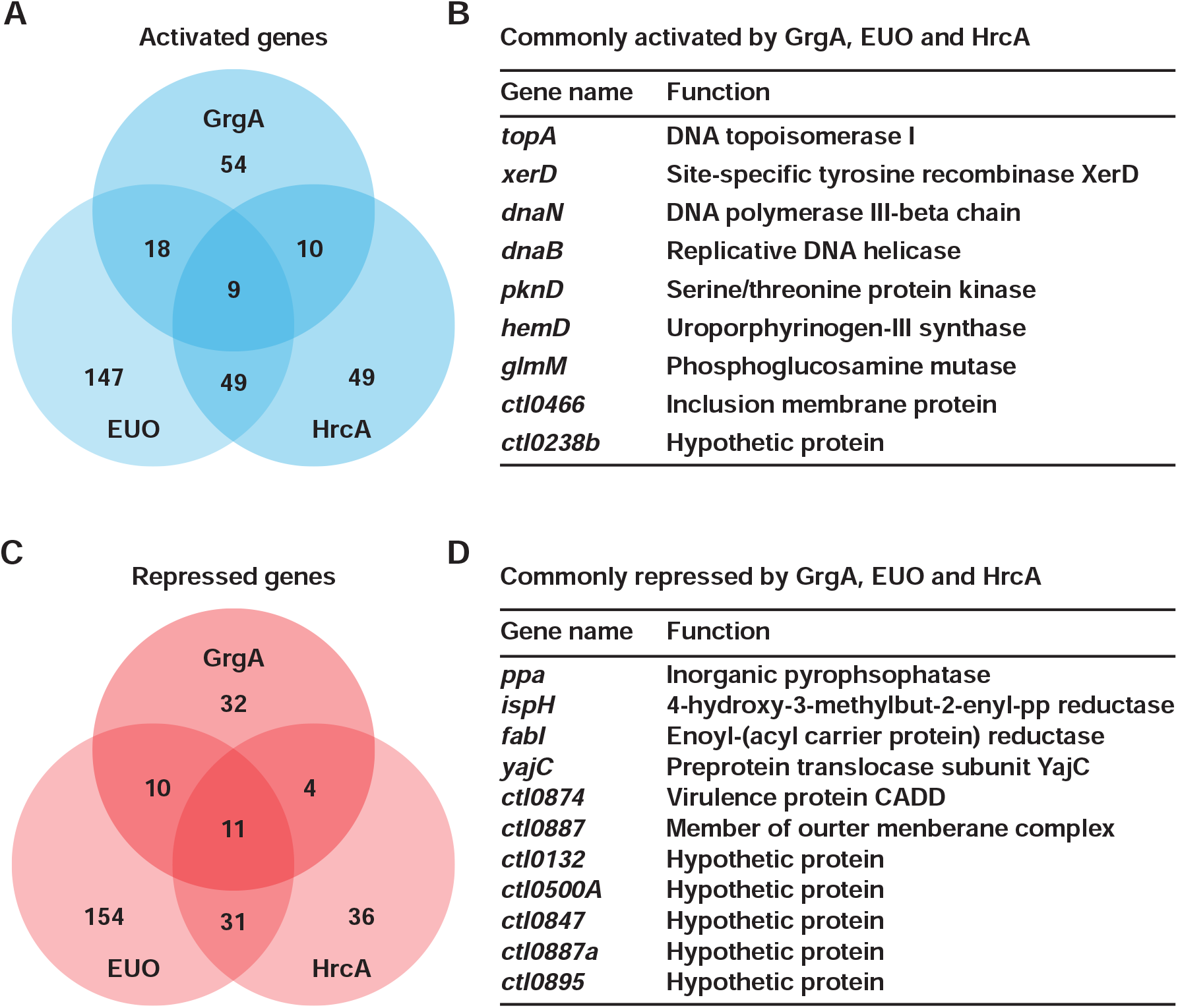
GrgA-, Euo-, and HrcA-coregulated genes. (A, C) Venn diagrams showing numbers of genes activated (A) and repressed (C) by overexpression of each of the 3 TFs. (B, D) Lists of genes commonly activated (B) and repressed (D) by GrgA, Euo, and HrcA overexpression. Note that genes commonly activated and repressed by any two of the TFs are shown in Table S15 and S16, respectively.

### GrgA-controlled transcriptional network

Using RNA-seq data (Tables S10) and RT-qPCR data (Figs. 6, S9) obtained from GrgA transformants, we elucidated a GrgA-regulated transcriptional network 30 min, 1 h, and 2 h after ATC treatment. Within 30 minutes of ATC treatment, GrgA activated expression of 12 molecules including the TFs Euo and HrcA, and repressed expression of two genes (*trpB* and *ctl0887*) (Fig. 10A). As will be discussed, the repression is likely an indirect effect of GrgA overexpression.

**Fig. 10.**
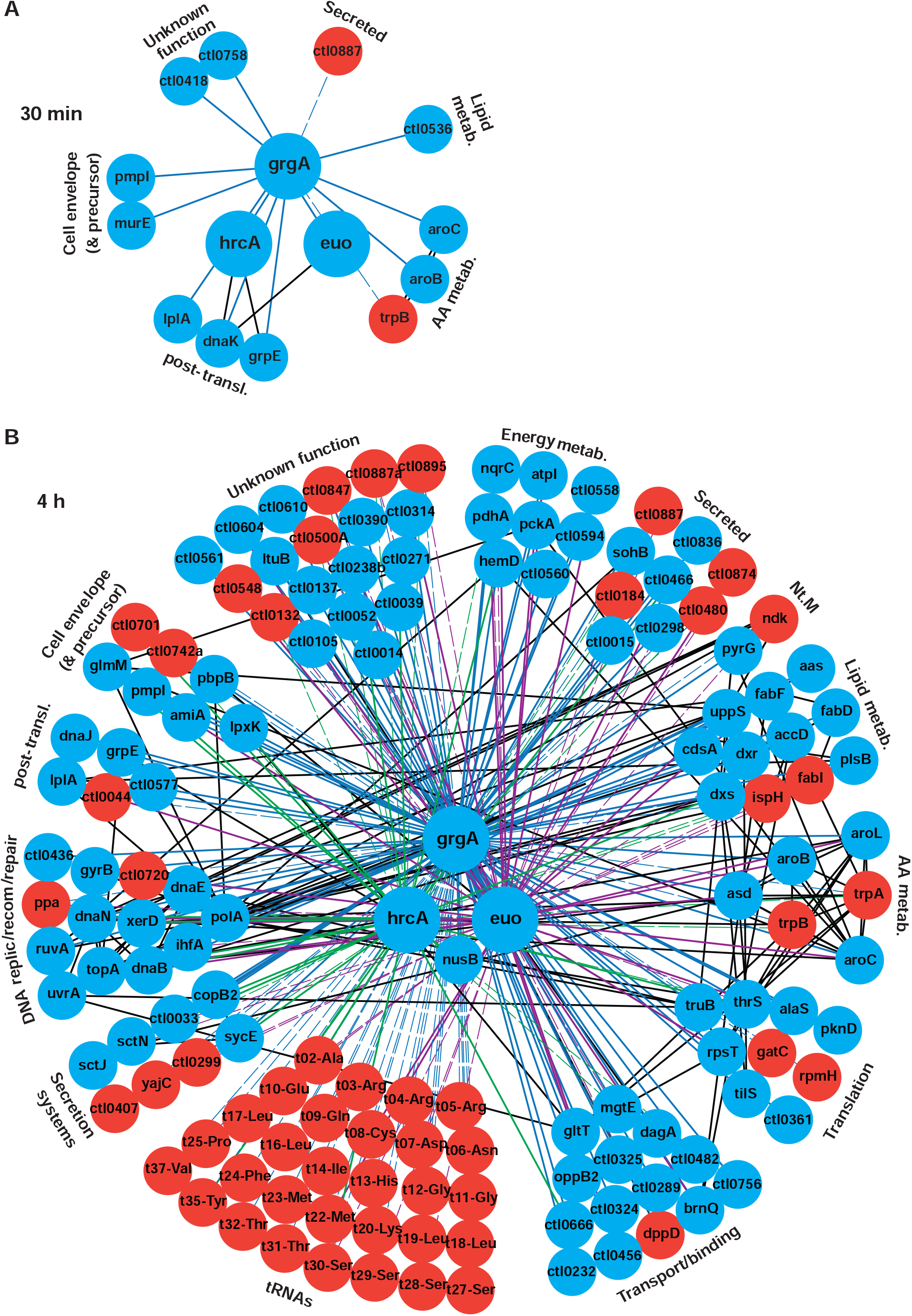
GrgA-regulated transcriptional network. (A) Network established within 30 min of ATC-induced GrgA overexpression. (B) Network developed by 4 h of ATC-induced GrgA overexpres-sion. (A, B) Blue and red nodes are genes activated and repressed by GrgA, respectively. Solid and dashed lines connect TFs to activated and repressed genes, respectively. Blue, purple and green lines connect GrgA, Euo, and HrcA, respectively, to their target genes. Black lines connect genes in correlation identified by STRING-db v.11. Functional group are labeled. Abbreviations: AA, amino acid; metab., metabolism; Nt.M, nucleotide metabolism; Post-transl., posttranslational protein modification; recom, recombination; replic, replication.

Large numbers of additional genes were activated and repressed at 1 h and 2 h after ATC treatment in GrgA transformants. The products of these genes can be classified into at least 16 clusters (Fig. S12). Overall, the networks at these two points are similar with only one major difference: 14 versus 4 downregulated tRNAs. Given the fact that 30 tRNA are downregulated at 4 h (Fig. 10B), we suspect that our RNA-seq samples for these two time points have inadvertently been switched around.

We developed the 4 h network by analyzing RNA-seq data obtained from not only GrgA transformants (Table S5), but also Euo and HrcA transformants (Tables 13, 14) to show how Euo and/or HrcA may mediate some of the transcriptomic changes in GrgA transformants induced with ATC. A still version of the network is presented in Fig. 10B, whereas an interactive version is provided as online supporting information (Fig. S13). The most striking event in this network is the downregulation of 30 tRNAs. Of these 30 tRNAs in GrgA transformants treated with ATC, 6 were also downregulated in the Euo transformants treated with ATC, suggesting the possibility that Euo may mediate tRNA expression in response to GrgA. However, 2 tRNAs downregulated by GrgA overexpression were also upregulated by Euo overexpression.

Other than tRNAs, there are slightly more genes in gene categories whose expression were changed by GrgA overexpression, compared with the networks developed for 1 h and 2 h (Fig. S13). RNA-seq data is consistent with the above-stated notion that while activating both Euo and HrcA expression, GrgA is repressed by both Euo and HrcA. Euo is in turn also repressed by HrcA. However, HrcA mRNA is significantly increased in Euo transformants treated with ATC for 4 h, suggesting that the long-term overall effect of Euo overexpression leads to increased HrcA expression even though it briefly downregulated HrcA expression at 15 min (Fig. S12B).

To recapitulate, our results presented above reveal pathways through which GrgA overexpression causes chlamydial growth inhibition via Euo- and HrcA-dependent and independent transcriptomic modulation.

## DISCUSSION

In this report, we determined the effects of GrgA overexpression on chlamydial growth and transcriptomic expression through experimentation with full-length GrgA and GrgA deletion mutants. We identified direct and indirect regulons of GrgA, and uncovered a TRN that encompasses GrgA, Euo, and HrcA. We further documented the inhibitory effects of Euo and HrcA overexpression on chlamydial growth and transcriptomic expression. Our findings have important implications for progression of the chlamydial developmental cycle.

### The direct and indirect regulons of GrgA

Our RNA-seq and RT-qPCR analyses revealed the direct and indirect regulons of GrgA by detecting time-dependent transcriptomic changes following ATC-induced GrgA overexpression. The direct regulon includes 12 genes that are activated within 10 to 30 min of ATC treatment (Figs. 6, S9; Tables S10, S11). mRNAs of the *C. trachomatis* L2b strain (a variant of CtL2) have an average half-life of only 15 min (43). While this short half-life suggests 10 to 30 min of ATC treatment is adequate to identify most activated and repressed genes, a longer treatment duration may be required to detect changes in RNAs with extraordinarily long half-lives. Therefore, it is possible that we have not identified all direct targets of GrgA.

Several lines of evidence presented in this report affirm that GrgA activates both σ^66^- and σ^28^-dependent promoters through direct interactions with σ^66^ and σ^28^, a notion drawn from our previous *in vitro* studies (35–37). Overexpression of the σ^66^-binding-defective Δ1-64 GrgA did not affect chlamydial growth, whereas full-length GrgA overexpression induced severe growth inhibition. Overexpression of the σ^28^-binding-defective Δ138-165 GrgA caused mild growth inhibition, albeit by a magnitude much less than that caused by full-length GrgA overexpression. Additionally, full-length GrgA overexpression caused numerous transcriptomic changes while Δ1-64 GrgA overexpression caused only a small increase in PmpI mRNA. Furthermore, among 8 promoter regions upstream of the 4 single-gene units and 4 operons activated within 10 to 30 min of ATC induction of GrgA, 5 have conserved σ^66^-dependent promoter elements and 1 has recognizable σ^28^-dependent promoter elements (Fig. 6, S10). Finally, the transcription activities of two of the σ^66^-dependent promoters (Euo and HrcA) and the σ^28^-dependent *pmpI* promoter are stimulated by GrgA *in vitro*.

Two genes (*trpB* and *ctl0887*) are downregulated 30 min after ATC treatment (Fig. 10 & Tables S10, S11). These genes are likely subjects to indirect rather than direct repression by GrgA. Neighboring genes encoded by different DNA strands can be activated or repressed by a bacterial TF if they share an intergenic promotor region (44). Because *trpB* and *ctl0887* are located far from any GrgA-activated genes, the opportunity for direct repression is unlikely. Indeed, RNA-seq data from HrcA and Euo transformants suggest that GrgA downregulates *trpB* expression indirectly through HrcA, and downregulates *ctl0887* expression through Euo and/or HrcA. Nonetheless, we cannot completely rule out the possibility that GrgA acts as a repressor since dual functional TFs have been identified in other bacteria (45).

Excluding the 12 direct target genes induced by 30 min, we consider all genes whose RNA levels significantly increased or decreased between 1 h and 4 h as constituents of the indirect regulon of GrgA. Because GrgA is an activator of *euo* and *hrcA*, it is not surprising that the indirect regulon is much larger than the direct regulon. Nearly 150 target genes were identified by 4 h when an arbitrary threshold of 33% change was applied. This approximation likely underestimates the true size of the regulon. As discussed above, physiological targets that show a percent change less than 33% are excluded. Furthermore, the fact that GrgA overexpression results in reduced RB replication suggests that the numbers of RNA reads should be normalized with the genome copy for cultures subjected to long ATC treatment (e.g., 2 to 4 h). However, there is no proper way to perform normalization because the RNA samples undergo procedures of host and bacterial rRNA removal and polyadenylated mRNA removal before library construction (47). Thus, we likely have underestimated the number of activated genes and the degrees of their activation, and at the same time overestimated the number of downregulated genes and the degrees of their downregulation.

### Chlamydial growth and development controlled by GrgA-directed TRN

Activation of *euo* and *hrcA* transcription by GrgA and the expression profiles of the three TFs indicate that GrgA serves multiple roles in the chlamydial cycle. *Euo* was initially identified as an immediate early gene in *Chlamydia psttaci* (33). Microarray studies confirmed that *euo* is immediately transcribed *C. trachomatis* EBs enter host cells (8, 9). Microarray detected GrgA mRNA from 8 hpi through 40 hpi (8). Our own expression analysis using western blotting (35) and work by Skipp et al. using quantitative proteomics (48) both detected high levels of GrgA in both EBs and RBs. Furthermore, protein mass spectrometry carried out by Saka et al. also detected GrgA in EBs although they failed to observe GrgA in RBs (49). We speculate that the GrgA protein prepacked into EBs plays a critical role in activation of *euo* transcription immediately following host cell entry. The Euo protein functions as a master repressor of late genes (11–13). By binding to the promoters of late genes and repressing their expression, EBs can utilize limited resources to express early genes required for converting themselves into proliferative RBs.

We believe that GrgA is also a physiological activator of *hrcA* transcription starting 24 hpi (8, 9). HrcA is known as a heat-inducible TF in bacteria (50). However, *C. trachomatis* infection seldom induces fever in infected humans. Its cyclic expression takes place *C. trachomatis* cultured at 37 °C (8, 9), a point at which RBs start to convert back to infectious EBs. Thus, HrcA either plays an active role in the redifferentiation by repressing its target genes or by keeping them silent in EBs. Consistent with this interpretation, there are examples that HrcA controls cell cycle-dependent protein expression in bacteria at normal growth temperature and plays only a minor role in heat shock response (51, 52).

A question arises as to why GrgA would not activate *hrcA* when it activates *euo*. GrgA could have differential affinity for the promoters of *euo* and *hrcA in vivo*, as promoter hierarchy is common among TF regulons (44). In addition, the chromatin configuration, which can significantly influence transcription, differs drastically in EBs and RBs.

How GrgA regulates RB growth during the midcycle is less obvious. Given the large number of genes in its indirect regulon (Fig. 10), GrgA likely fulfils its function as a growth regulator through balanced action of its direct and numerous indirect target genes with roles in biosynthesis, metabolism and other processes. Similarly, many genes (e.g., tRNA genes) regulated by GrgA may coordinate the transition of RBs to EBs during late developmental stages.

We identified 9 commonly activated and 11 commonly repressed genes in GrgA, Euo, and HrcA transformants undergoing growth arrest due to ATC-induced overexpression of respective TFs (Fig. 9; Tables S15, S16). Their disregulated expression may contribute to or result from chlamydial growth inhibition. Paradoxically, 4 of the 9 commonly activated genes encode proteins involved in DNA replication and repair, which include topoisomerase I, DNA polymerase III, DNA helicase, and a site-specific tyrosine recombinase (XerD). Interestingly, all these four genes are also upregulated during interferon-γ-induced chlamydial persistence when growth is also reduced. In addition, Euo and mRNA of CT505, transcription partner of GrgA, are increased under the condition (53). Thus, GrgA and Euo likely regulate chlamydial persistence, which can be induced by cytokines and antibiotic treatment (53–55).

The fifth commonly activated gene *pknD* encodes a protein kinase (56, 57). It is known that inhibition of either PknD or a chlamydial phosphatase PP2C that dephosphorylates PknD can impede chlamydial growth (58, 59). Given the importance of the balance between protein phosphorylation and dephosphorylation, increased PknD expression likely contributes to chlamydial growth inhibition. It is not apparent how increased expression of four remaining commonly activated genes, *hemD, gluM, ctl 0466 and ctl0238b* contributes to growth inhibition.

6 of the 11 genes commonly repressed by GrgA, Euo, and HrcA overexpression encode functionally known proteins whose dysregulation may negatively affect chlamydial growth. The *ppa-*encoded enzyme is required for enterobacterial DNA replication (60). The enzyme encoded by *ispH* produces isopentyl diphosphate. The isoprenoid is a precursor of peptidoglycan (54, 61, 62). In addition to inhibited peptidoglycan synthesis, downregulated IspH may lead to isoprenoid precursor accumulation, which may alter chlamydial gene expression through regulating the interaction of histone with DNA (63, 64). The product of *fabI* is a key enzyme required for chlamydial growth that acts in the type II fatty acid synthesis system (65). The product of *yajC* is a constituent of the preprotein translocase, which is required for protein export across the inner membrane, an essential function in Gram-negative bacteria (66). CADD interacts with death receptors on the host cells (67, 68), and may facilitate EB release by inducing host cell death in a late developmental stage. Finally, CTL0887 is a member of the chlamydial outer membrane complex (69, 70). Although the exact function of CTL0887 remains unknown, the complex is required for maintaining the integrity of the bacterium.

Numerous tRNAs are downregulated following GrgA overexpression (Table S5; Fig. 10), which would contribute to growth inhibition. A smaller number of tRNAs are also downregulated following Euo overexpression (Fig. 10; Tables S13, S16). Other bacteria downregulate tRNA transcription in response to nutrient deprivation (71, 72). This stringent response phenomenon is mediated by (p)ppGpp, a protein produced only during starvation that acts directly on the RNAP holoenzyme to reprogram transcription. *Chlamydia* lacks the capacity to synthesize (p)ppGpp however (7, 41). It is likely that chlamydial tRNA expression is temporarily delayed in the immediate early stage and later downregulated as RBs converts to EBs.

In summary, we have identified at least 12 genes that are direct targets of GrgA, the newest transcription factor in *Chlamydia.* By activating expression of two major transcription factors, Euo and HrcA, and by regulating expression of numerous additional genes with functions in almost all cellular processes, GrgA acts as a master transcription regulator that controls chlamydial growth and development. It may also regulate chlamydial persistence, an important clinical phenomenon. Hopefully, an efficient gene-silencing technology not only applicable to nonessential genes but also essential genes will soon be developed to illuminate the precise roles of GrgA in chlamydial physiology (73).

## MATERIALS AND METHODS

### Plasmids

Plasmids used for this study are listed in Table S18. Primers used to amplify fragments are listed in Table S19. All primers were custom synthesized at Sigma. pGFP::SW2-GrgA (Fig. S1) was constructed by fusing a PCR-amplified full-length GrgA fragment with *Sal*I-cut pGFP::SW2 (38). The GrgA fragment was amplifying by *PfuUltra* DNA Polymerase (Agilent, Cat. # 600380). Fusion was performed with the Cold Fusion Cloning Kit (SBI System Biosciences, Cat. #MC010B-1).

pTRL2-NH-GrgA (Fig. S1) was constructed using the Cold Fusion Cloning Kit to combine two DNA fragments. Fragment 1, also termed fragment TRL2(Δgfp), was amplified by the *PfuUltra* DNA polymerase using pASK-GFP-L2-mkate2 (74) as the template. Fragment 2 encoded NH-GrgA and was amplified by the same enzyme with pET21a-NH-GrgA as the template. pTRL2-GrgAΔ1-64 and pTRL2-NH-GrgAΔ138-165 were constructed using the QuickChange II Site-directed Mutagenesis Kit to delete DNA sequences from a pTRL2-NH-GrgA template that encode amino acid regions 1-64 and 138-165.

Fragment amplification for constructing remaining plasmids was performed using Q5 high-fidelity DNA polymerase (NEB, Cat. # M0491). A TRL2(Δgfp)-NH fragment was amplified using pTRL2-NH-GrgA as template. The Euo and HrcA encoding fragment was amplified by using CtL2 genomic DNA as the template, and were fused to fragment TRL2(Δgfp)-NH using the NEBuilder HiFi DNA Assembly Cloning kit (NEBuilder, NEB, Cat. #M0491) to create plasmids pTRL2-NH-EUO and pTRL2-NH-HrcA, respectively.

An *euo* promoter fragment and an *hrcA* promoter fragment (Supplemental sDoc1) were amplified using CtL2 genomic DNA as templates, and were fused to vector fragments using NEBuilder to create pMT1125-Peuo and pMT1125-PhrcA, respectively. Promoter fragments were amplified using pMT1125 (75) as template.

pMT1125-PpmpI(σ^28^) and pMT1125-PpmpI(σ^66^) were constructed in two steps. First, the putative σ^28^-dependent and σ^66^-dependent promoter fragments (Supplemental sDoc1) were amplified using CtL2 genomic DNA as the templates. Resultant DNA fragments were digested with *Xba*I and *EcoR*V and ligated to *Xba*I/*EcoR*V-digested pMT1125 using T4 DNA ligase. Second, the quinine nucleotide inside the *EcoR*V site was deleted using Q5 site-directed mutagenesis kit (Cat. #E0554S).

Plasmids constructed were subject to Sanger sequencing at Genscript or Psomogen to ensure sequence authenticity. For chlamydial expression vectors, sequencing analysis also covered the CtL2-encoded genes and additional applicable elements (i.e., the Pnm promoter, EGFP-Cat gene, ATC-inducible promoter, and/or tet repressor-coding sequence), in addition to the coding sequences of TFs or their deletion mutants. Promoter fragments and the reporter cassette in pMT1125-derived vectors were sequenced.

#### CtL2 strains

Wild-type CtL2 (strain 434/BU) was purchased from ATCC (76). EBs were purified from L929 cells via MD-76 gradient ultracentrifugation as described previously (77). Titers of EB stocks were determined as follows. L929 cells grown on 96-well plates were infected by centrifugation for 20 minutes at 900 × g. These infected cells were then fixed with cold methanol at 30 hours post-inoculation (hpi) and stained successively with two antibodies: a monoclonal L21-5 anti-major outer membrane protein antibody (78) and an FITC-conjugated rabbit anti-mouse antibody (79). Transformation was performed as described (80) with modifications (81). 1.3 × 10^7^ IFUs of EBs were mixed with 4-6 μg of plasmid DNA in 50 μl CaCl_2_ buffer (10 mM Tris, pH 7.4 and 50 mM CaCl_2_) and incubated for 30 minutes at room temperature. The mixture was then diluted with 1.2 mL Hanks Balanced Salt Solution (HBSS; Sigma, Cat. # D8622) and used to inoculate a 6-well plate of nearly confluent L929 cells (i.e., ~0.2 ml of the suspension per well). Monolayers were infected at room temperature by centrifugation for 20 minutes at 900 × g, after which HBSS was replaced with DMEM containing 5% FBS (2 ml/well). Cultures were supplemented with cycloheximide (final concentrations: 1 μg/ml) and penicillin G (final concentration: 2 U/ml). Cells from each well were harvested into 500 μl HBSS at 36 hpi, disrupted by brief sonication and centrifuged for 10 min at 1000 × g and 4 °C. The supernatant was used thereafter to infect a new well of nearly confluent L929 monolayer on a 6-well plate by centrifugation. Immediately after infection, medium containing both cycloheximide and penicillin G was added, and the process of harvesting and infection was repeated. After RFP-expressing inclusions were noted (typically at the end of passage 2 or 3), the concentration of penicillin G was increased to 4 U/ml for the next passage and further to 10 U/ml for 2 additional passages. Thereafter, penicillin G was replaced with 10 μg/ml ampicillin for further expansion.

To generate transformant clonal populations, 6-well plates of L929 cells were inoculated with EBs (~1-6 inclusions/well) and cultured using medium with 20 μg/ml ampicillin. A P200 micropipette was used to sample one inclusion from each well at 24 to 28 hpi (i.e., 6 total inclusions were sampled from 6 different wells). Intracellular chlamydiae were released from each sampled inclusion by sonication for 5 seconds, centrifuged, and subsequently used to inoculate an entirely new 6-well plate of L929 cells. 6 additional inclusions were picked from a plate with observable inclusions (typically, only 2-3 of the 6 inoculated wells yielded inclusions). This process was repeated one more time to ensure homogeneity. For further experimentation, EBs of transformant clonal populations were prepared and purified as described above. Infectivity of EB stocks were determined as described, except that RFP-expressing inclusions of CtL2 transformants in live cultures were scored without immunostaining (73).

#### Determination of *C. trachomatis* growth

Nearly confluent L929 cell monolayers grown on 24-well (for determining progeny EB production) and 6-well (for microscopic analysis) plates were infected with MD-76 gradient-purified CtL2 transformants at a multiplicity of infection (MOI) of 1 IFU per 3 cells. Unless otherwise indicated, expression of GrgA, Euo, and/or HrcA was induced by replacing culture media with fresh media containing 10 nM ATC. To quantify progeny EB production, cells were harvested in 500 μL SPG buffer at 30 hpi; recoverable IFUs were determined as previously described (73). RFP-expressing inclusions were imaged at 36 hpi. The Java-based ImageJ software was then used to process the images. An empiric threshold value was first determined and applied, after which noise reduction and binarization calculations were performed. The analyze particles function was then called with minimum size and circularity constraints to compute potential inclusion boundaries within the given image. Visual inspection was conducted to ensure accurate particle identification and selection for subsequent intensity measurements.

#### Epi-fluorescence microscopy

L929 cell monolayers grown on 6-well plates were infected with EBs of CtL2 transformants at an MOI of 1 IFU per 3 cells and incubated with or without 10 nM ATC for 34 to 36 hours. Bright field and red fluorescent images were acquired on an Olympus IX51 fluorescence microscope using a constant exposure time for each channel. Image overlay was performed using the PictureFrame software.

#### Confocal fluorescent microscopy

L929 cell monolayers grown on coverslips were infected with EBs of NH-GrgA transformants at an MOI of 1 IFU per 5 cells. GrgA expression was induced with 20 nM ATC at 8 hpi. 6 h later, cells were rinsed with PBS, fixed by incubation in PBS containing 3% formaldehyde, 0.045% glutaraldehyde for 10 min, washed twice with PBS and permeabilized with 90% cold methanol (82). Chlamydiae were stained with polyclonal a rabbit anti-tRFP antibody (Evrogen, Cat. # 233), which recognized the RFP mKate protein, and then FITC-conjugated goat anti-rabbit IgG secondary antibody (Immunotech). Host cell cytoplasm was stained with 0.01% Evans Blue; host cell chromosomal DNA was stained with 1μg/mL Hoechst 33342. Cells were imaged using a Zeiss LSM710 confocal microscope equipped with a 100X Plan-Apochromat oil immersion lens.

#### Electron Microscopy

L929 cell monolayers were infected with GrgA transformants at an MOI of 1 IFU per cell. Cultures were then treated with or without aTC, collected in PBS containing 10% FBS at 14 hpi, and centrifuged for 10 minutes at 500 × g. Pelleted cells were resuspended in EM fixation buffer (2.5% glutaraldehyde, 4% paraformaldehyde, 0.1 M cacodylate buffer) at RT, allowed to incubate for 2 hours, and stored at 4 °C overnight. To prepare samples for imaging, cells were first rinsed in 0.1 M cacodylate buffer, dehydrated in a graded series of ethanol, and then embedded in Eponate 812 resin at 68 °C overnight. 90 nm thin sections were cut on a Leica UC6 microtome and picked up on a copper grid. Grids were stained with Uranyl acetate followed by Lead Citrate. TIFF images were acquired on Philips CM12 electron microscope at 80 Kv using AMT XR111 digital camera. RB diameters were measured using ImageJ software (83).

#### Cellular genomic DNA and RNA isolation

Total host and chlamydial genomic DNA and RNA were isolated from non-infected and chlamydia-infected L929 cells using TRI reagent (Sigma, Cat. # 93289), which separates DNA and RNA into different phases. DNA and RNA were purified in accordance with the manufacturer’s instructions (84). Genomic DNA was dissolved in a buffer containing 0.1 M HEPES and 8 mM NaOH. These samples were stored at −20 °C. RNA was dissolved in DEPC-treated H_2_O and further treated with RNase-free DNaseI to eliminate residual DNA contamination. The resultant DNA-free RNA samples were stored at −80 °C.

#### Quantitative PCR (qPCR) and reverse transcription qPCR (RT-qPCR)

Thermo Fisher QS5 qPCR machine was used for qPCR and RT-qPCR analyses to quantify relative CtL2 genome copy numbers and mRNA levels, respectively. Genomic qPCR was performed using Applied Biosystems PowerUp SYBR Green Master Mix (Thermo Fisher Scientific, Cat. # A25742) following manufacturer’s instructions. For each reaction, 5 ng of purified total host and bacterial genomic DNA was used as template. The primer pair were qPCR-ctl0631-F and qPCR-ctl0631-R (Table S10). RT-qPCR was performed using Luna Universal One-Step RT-qPCR kit (NEB, Cat. # E3005E) following manufacturer’s instructions. For each reaction, 600 ng of purified total host and bacterial RNA was used as initial template for cDNA synthesis. All genomic and RT-qPCR reactions were performed in technical duplicate or triplicate.

#### RNA sequencing

Total RNA integrity was determined using Fragment Analyzer (Agilent) prior to RNA-seq library preparation. Illumina MRZE706 Ribo-Zero Gold Epidemiology rRNA Removal kit was used to remove mouse and chlamydial rRNAs. Oligo(dT) beads were used to remove mouse mRNA. RNA-seq libraries were prepared using Illumina TruSeq stranded mRNA-seq sample preparation protocol, subjected to quantification process, pooled for cBot amplification and sequenced with Illumina HiSeq 3000 platform with 50 bp single-read sequencing module. In average, 20-25 million reads were obtained for each RNA-seq sample. Short read sequences were first aligned to the CtL2 chromosome (accession # NC_010287.1) and the transformed plasmids using TopHat2 aligner and then quantified for gene expression by HTSeq to obtain raw read counts per gene, and then converted to RPKM (Read Per Kilobase of gene length per Million reads of the library) (85–87).

### TRN development

Pathway analysis was first performed on significantly regulated gene sets whose P values were < 0.05 by STRING-db v.11 and modified to increase font size, nodes and edges were changed by color coding. Secondly, add more GrgA-regulated genes pathway (edges) according RNA sequencing data without altering original network relationships.

### *In vitro* transcription assay

Chlamydial RNA polymerase holoenzyme was partially purified from RBs of pTRL2Δgfp-transformed CtL2 using Heparin Agarose (Sigma) as previously described (35). *In vitro* transcription assays for σ^66^-dependent promoters and σ^28^-dependent promoters were performed as previously described (35, 36).

### Western Blotting

L929 cells grown on 6-well plates were infected with transformants. Expression induction was performed at 14 hpi using 10 nM ATC. Cells were harvested in 100 μL 1X SDS-PAGE sample buffer at 15 hpi, heated at 95 °C for 5 min, and sonicated for 1 minute at 35% amplitude (5 second on, 5 seconds off). Proteins were resolved in 10% SDS-PAGE gels and transferred onto PVDF membranes. GrgA and mutants were detected using a monoclonal anti-His antibody (Genscript, Cat. A00186) and a mouse anti-GrgA antibody (35).

### Statistical analysis

R package DESeq was used to normalize data and find group-pairwise differential gene expression based on three criteria: Pval < 0.05, average rpkm > 1, and fold change ≥ 1. Genes were clustered into groups based on temporal patterns of transcriptomics using Gaussian mixture models (88). All other quantitative data were analyzed using *t* tests in Excel of Microsoft Office.

## ACKNOWLEDGEMENTS

We thank Dr. Huaye Zhang for assistance with the confocal microscopy, Mr. Rajesh Patel for assistance with electron microscopy, Dr. Joseph Fondell for discussions. We also thank Dr. P. Scott Hefty (University of Kansas) for the supply of pASK-GFP/mKate2-L2. This work was supported by grants from the National Institutes of Health (grant # AI122034 and AI140167 to HF) and New Jersey Health Foundation (grant # PC20-18 to HF). Genome Sequencing Facility at UTHSA is supported by NIH-NCI P30 CA054174 (Cancer Center at UT Health San Antonio), NIH Shared Instrument grant 1S10OD021805-01 (S10 grant), and CPRIT Core Facility Award (RP160732).

## Notes

### Competing Interest Statement

The authors have declared no competing interest.

